# Spatial Tracking Across Time (*STAT*): Tracking Neurons Across In-Vivo Imaging Sessions through Optimizing Local Neighborhood Motion Consistency

**DOI:** 10.1101/2023.05.13.540658

**Authors:** Shijie Gu, Emily L. Mackevicius, Michale S. Fee, Pengcheng Zhou

**Affiliations:** McGovern Institute, Department of Brain and Cognitive Sciences, Massachusetts Institute of Technology, Cambridge, MA, USA; University of California Berkeley and University of California San Francisco Joint Program in Bioengineering, San Francisco, CA, USA; Columbia University, New York, NY, USA; Faculty of Life and Health Sciences and the Brain Cognition and Brain Disease Institute, Shenzhen Institute of Advanced Technology, Chinese Academy of Sciences, Shenzhen, China

## Abstract

Chronic calcium imaging has become a powerful and indispensable tool for analyzing the long-term stability and plasticity of neuronal activity. One crucial step of the data processing pipeline is to register individual neurons across imaging sessions, which usually extend over a few days or even months, and show various levels of spatial deformation of the imaged field of view (FOV). Previous solutions align FOVs of all sessions first and then register the same neurons according to their shapes and locations [1, 2]. However, the FOV registration is computational intensive, especially in the case of nonrigid case.

Here we propose a cell tracking method that does not require FOV image registration. Specifically, the algorithm ***STAT*** (short for ***S***tay ***T*** ogether, ***A***lign ***T***ogether, and for ***S***patial ***T***racking ***A***cross ***T***ime) represents neurons from two sessions as two sets of neuronal centroids, uses point set registration (PSR) to find a spatially smooth transformation to align them while assigning correspondences. The optimization method iteratively updates between the general motion and individual neuron identity tracking, an idea seen in the computer vision literatures [3, 4]. Our method can be thought of as a specialization and simplification of these more general methods to calcium imaging neuron tracking.

We validate STAT on datasets with simulated nonrigid motion that is hard to motion correct without extensive manual intervention. Next, we test STAT on experimental data from singing birds collected on three different days, and observe stable song-locked activity across days. An example use case of this package is reference [5].

## 1 The Problem and the Intuition

Functional calcium imaging is a powerful method to measure the activity of individual neurons in behaving animals in real time. Populations of neurons in the brain can be genetically manipulated to express calcium-sensitive fluorescent indicators, such that when neurons become active, calcium influx into the cell body or other neuronal processes leads to large changes in fluorescence. Together with new imaging technologies, such as two-photon microscopy and head-mounted fluorescence microscopes, this approach allows the simultaneous recording of many tens or even thousands of neurons [6]. In some cases, these recordings can be carried out over many days to track neuronal activity under different conditions or across time as the animal learns a new task or behavior [7, 8]. A key challenge has been to analyze the functional imaging data to extract and quantify the underlying neural activity. In recent years effective methods have become available for extracting neural signal from raw calcium imaging videos [1, 2, 9, 10], but tracking these signals from individual neurons across multiple days is remains a challenge [1, 2].

The fundamental problem can be described as follows: Suppose we have two calcium imaging sessions and between the two the FOVs do not look exactly the same, potentially due to the vascular changes [11] that happen during the long non-imaging time between the two imaging sessions. Because within each imaging session, there is little motion, we could apply cell extraction algorithms to data from each day separately. The problem is to stitch together these two separate sets of neurons obtained from cell extraction algorithms: we want to know which neurons in session 1 correspond to which neurons in session 2. In figure 1, we show an example from our songbird data.

**Figure 1:**
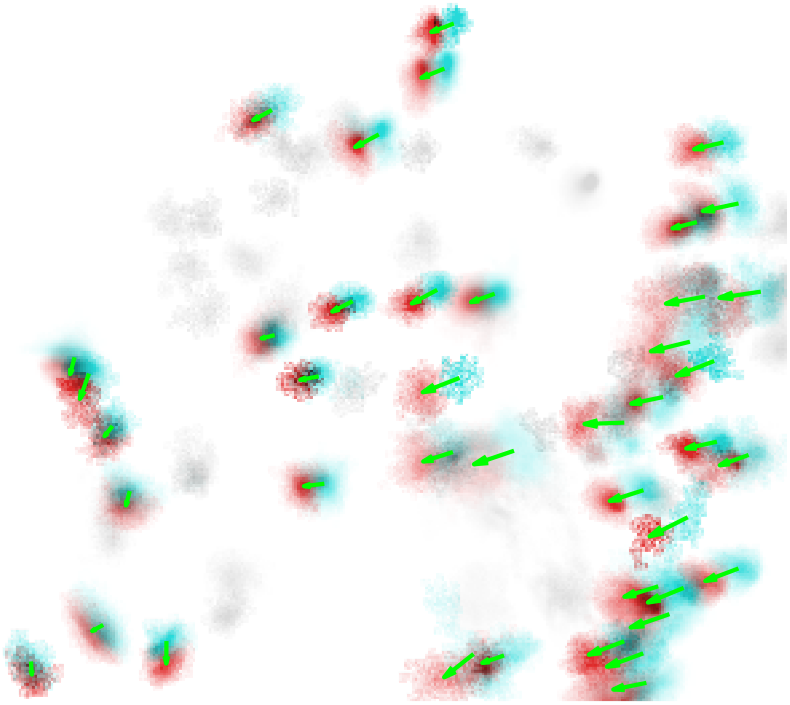
The challenge of tracking neurons: an example from real data. Spatial footprints of neurons imaged in singing juvenile zebra finch in two sessions, on two successive days. Neuron identities were hand-labeled in each session. Note that most neurons persisted across both sessions (cyan, session 1; red, session 2) but some neurons from session 1 did not appear in session 2, and vice-versa (gray). Neurons that were present in both sessions appear shifted in directions that are not identical across the entire field of view (green arrows indicate shift from session 1 to 2). Notably, however, nearby neurons appear to be shifted in similar directions by similar magnitudes.

In the example in figure 1 above, we see that:

- Motion is not the same across the whole field of view. In other words, motion is nonrigid. At the same time, nearby neurons move similarly [12].
- Even with 1-photon imaging, neuron footprints have some shape/orientation information. The same neuron has similar shapes across sessions [1].
- Each session does not have the same set of neurons. In other words, some neurons are only present in one session.

This paper proposes a method to track neurons, with the two considerations above, but we start describing our problem with the simplest case.

Suppose both session 1 and session 2 have the same set of *N* neurons, but just in different labels in each session. Then, what we are after is a permutation *i*_1→2_ that assigns a neuron in session 2 to each neuron *i* ∈ {1, 2, 3…*N*} in session 1. Alternatively we can express the correspondences using an assignment matrix *W* of size *N* by *N*, with 1’s in the entry *w*_*ij*_ if neuron *i* in session 1 corresponds to neuron *j* in session 2, and zeros otherwise. It is easy to see that in this case each row or each column of *W* sums to 1.

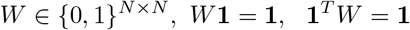

where the notation “**1**” means a column vector of 1’s and “*X*^*T*^” means taking the transpose of the matrix *X*.

In real data, however, one session tends to have some unique neurons that do not appear on the other session. Thus in a more general sense, if session 1 has *n*_1_ neurons and session 2 has *n*_2_, we are looking for a sparse binary **assignment matrix** *W* that is of size *n*_1_ by *n*_2_, with 1’s in the entry *w*_*ij*_ if neuron *i* in session 1 corresponds to neuron *j* in session 2, and zeros otherwise. Because now each row or each column of *W* can be all zeros (no-match case), we also revise our constraint on *W* so that each row sum or column sum is at most 1:

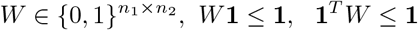

To find this assignment matrix, we use the observations above about the smooth mapping and neuronal shape. We formalize the intuitions above together into an optimization problem. Shape information can be used for initialization of the assignment of the match matrix, and smooth mapping is formulated into a cost function, through minimizing which we update assignments in iterations in a way. The convergence to the minimum of the cost function results in a substantial improvement in matching performance compared to the initialization.

Below, we give an overview of the algorithm, followed by definitions of variables that are essential in the algorithm. Then, we describe our algorithm in detail.

## 2 Algorithm Overview

Figure 2 gives an overview of our algorithm (variable notations shown on the figure will be defined in the next section). First, during optimization, we only use cell **centers** rather than the whole cell shape. This allows us to work with low dimensional variables in the optimization. Figure 2 (A) and (B) showed the centers of neurons in session 1 (cyan) and session 2 (red) in 3D perspective. Figure 2 (C) shows the projection of neurons from session 1 onto session 2. Figure 2 (D) then initializes the assignment, the result of which is shown in dashed arrows. The arrows are also the **displacement vectors** of each neuron in session 1. From there, the algorithm starts to update the assignment by minimizing neighboring **motion discrepancy**. Take the cyan neuron in the upper left corner as an example: it was assigned to the red neuron above initially. However, because other neighboring neurons are all assigned to neurons on the right, the updating process would then assign the cyan neuron in consideration to the red neuron to its right so that neighboring motion discrepancy is minimized. In this process, we use a **threshold** so that the only assignments that give a discrepancy below the threshold are allowed. This threshold is essentially the way our algorithm detects unique neurons as outliers. Take the cyan neuron in the upper center of the FOV as an example: because all the potential assignments would all give a big motion discrepancy with others, the final output (figure 2E) does not include an assignment for this neuron, i.e., it successfully detects the neuron as a neuron unique to one session.

**Figure 2:**
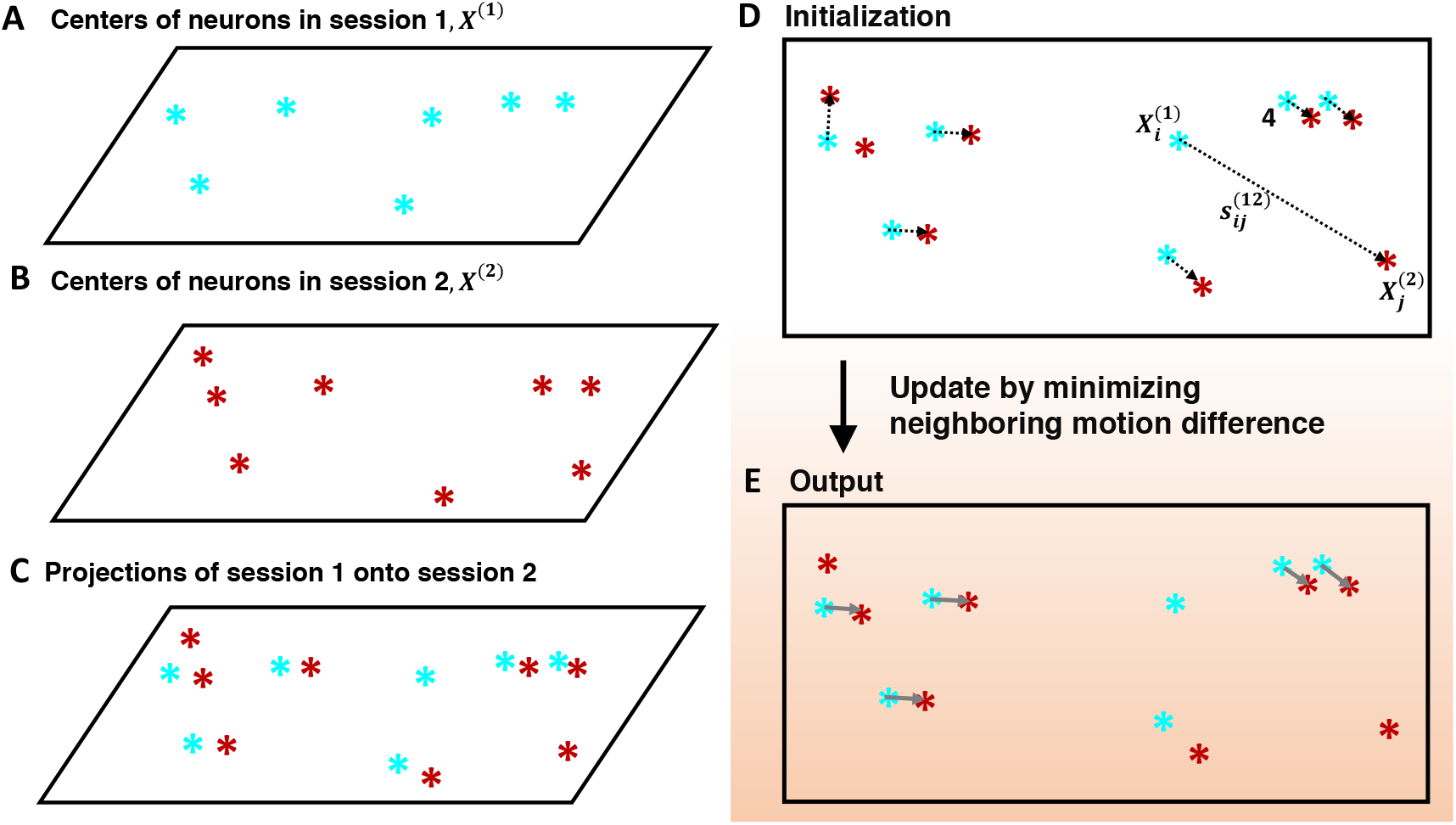
*STAT* overview.

The bold phrases in the paragraph above include the most important variables in the algorithm, which we now define with mathematical notations in the next section.

## 3 Variable Definitions

1. Neuron spatial footprints on session 1 and session 2: Because our algorithm is downstream of cell extraction analysis, we follow a similar notation used in a few calcium imaging cell extraction methods [9, 13] and refer to “neuron spatial footprint” as the neuron shape in a specific location in the FOV.

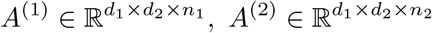

where *d*_1_ and *d*_2_ are the number of row, column pixels in each frame, and *n*_1_ and *n*_2_ are the number of neurons extracted in session 1 and session 2 respectively. In the 3D case, we have

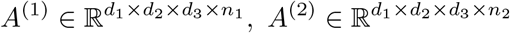

where *d*_3_ is the pixel number in the third dimension. The spatial footprint of each neuron has two pieces of information about each neuron: (1) the center of mass of the spatial footprint (2) the geometric shape of each neuron. Our algorithm could use the shape of neurons to initialize, but we use neurons’ centers as the primary piece of information.
2. Neuron centers on sessions 1 and 2:

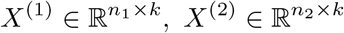

where *k* is the dimension of FOV. For single plane imaging data *k* = 2, and for volume imaging data *k* = 3. Further, for a specific neuron *i* in session *a*, we refer to its coordinate on session *a* as 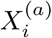, for *i* ∈ 1 ≤ i ≤ *n*_*a*_. This notation is defined similarly in the case of 3D volume data.
3. The displacement vector: The essence of our algorithm is to minimize motion discrepancy. To simplify notation, We first define the displacement vector. The displacement vector between neuron *i* in session 1 and neuron *j* in session 2 is:

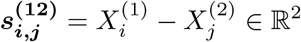

We similarly define displacement vectors between any pair of neurons within session 1 and session 2:

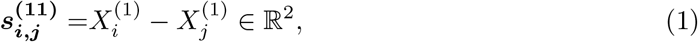

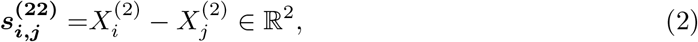
4. Assignment matrix: We have already defined the assignment matrix in the last section, but because the matrix is sparse, we here also define the list form of the assignment matrix. *i*_1→2_ = 0 stands for the case where there is no neuron in session 2 assigned to it as all entries in row *i* of *W* are zeros. This no-match case is similarly defined for *j*_1*←*2_ = 0.

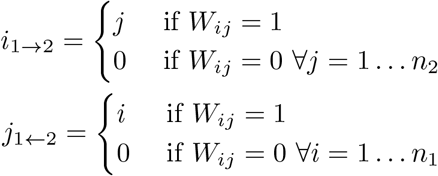 Note that {(*i, i*_1→2_)} or {(*j*_1*←*2_, *j*)} are equivalent to *W*. (*i, i*_1→2_), (*j*_1*←*2_, *j*) are useful in notating specific pairs of neurons. As an example, for a pair of matched neurons (*i, i*_1→2_), its displacement vector is 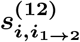. When *i*_1→2_ = 0, i.e., the neuron *i* has no match, 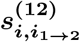 does not exist.
5. Motion discrepancy: We call 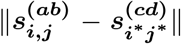 the discrepancy of motion vectors between neuron *i* in session *a* onto neuron *j* in session *b* and neuron *i*^*∗*^ in session *c* onto neuron *j*^*∗*^ in session *d*, where *a, b, c, d* ∈ {1, 2}.

## 4 Problem Formulation and Algorithm Details

We introduced in the last section the discrepancy of motion vectors between each neuron and its neighbors. The weighted version of the discrepancy is exactly the cost function in our algorithm, which we minimize in a greedy way to refine the assignment matrix. In this section, we describe in the detail the algorithm.

### 4.1 The Discrepancy Cost

Suppose neuron *i* in session 1 is assigned to neuron *j* in session 2, then we define the **discrepancy cost** *m* as the weighted sum of motion discrepancy with neurons close to neuron *i*.

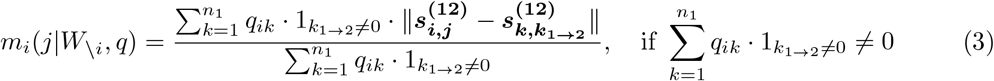

*W*_\*i*_ means looking at all other entries in *W* other than those in the *i*th row of *W*. The weight *q*_*ik*_ determines the importance of the neuron *k* when computing the motion discrepancy between the neuron *i* and the neuron *k* in session 1. The importance measure in *q*_*ik*_ can be defined flexibly. In our current implementation, we offer a few options and the simplest case is a step function, which sets *q*_*ik*_ = 1 if the distance between two neurons are smaller than *R* and *q*_*ik*_ = 0 if larger than *R*.

The way we define *m* makes it a good *confidence index*, because if an assignment has very similar motion to all other neighboring pairs, then very likely it would be a correct match. In other words, the smaller the discrepancy is, the more confident you are about the matching. (However, because of the error that is introduced in estimation of neuron centers, the relationship of *m* being smaller and assignment being more confident does not hold for extremely small values of *m*.)

To illustrate how the cost function works, we have in figure 3 an example. In particular, for the neuron 1, we demonstrate the calculation of *m*_1_(*j* |*W*_\*i*_, *q*), *j* ∈ {0, 1, 2, …, *n*_2_}. First, in figure 3(A), we use the simplest step function method to set *q*. In this situation, only neuron 2 and 3 would contribute to cost function *m*_1_(*j*|*W*_\*i*_, *q*), *j* ∈ {0, 1, 2, …, *n*_2_}. In addition, because neuron 2 and 3 both have their respective assignment, *k*_1→2_ = 0 for *k* = 2, 3, there are two components in the sum of the numerator of the equation 3 and the denominator becomes 1+1=2. Figure 3(B) explicates *m*_1_(*j*|*W*_*\i*_, *q*), *j* ∈ {0, 1, 2, …, *n*_2_}. We will explain why we *define* the case of *j* = 0. Other information in figure 3(B) will also be explained later.

**Figure 3:**
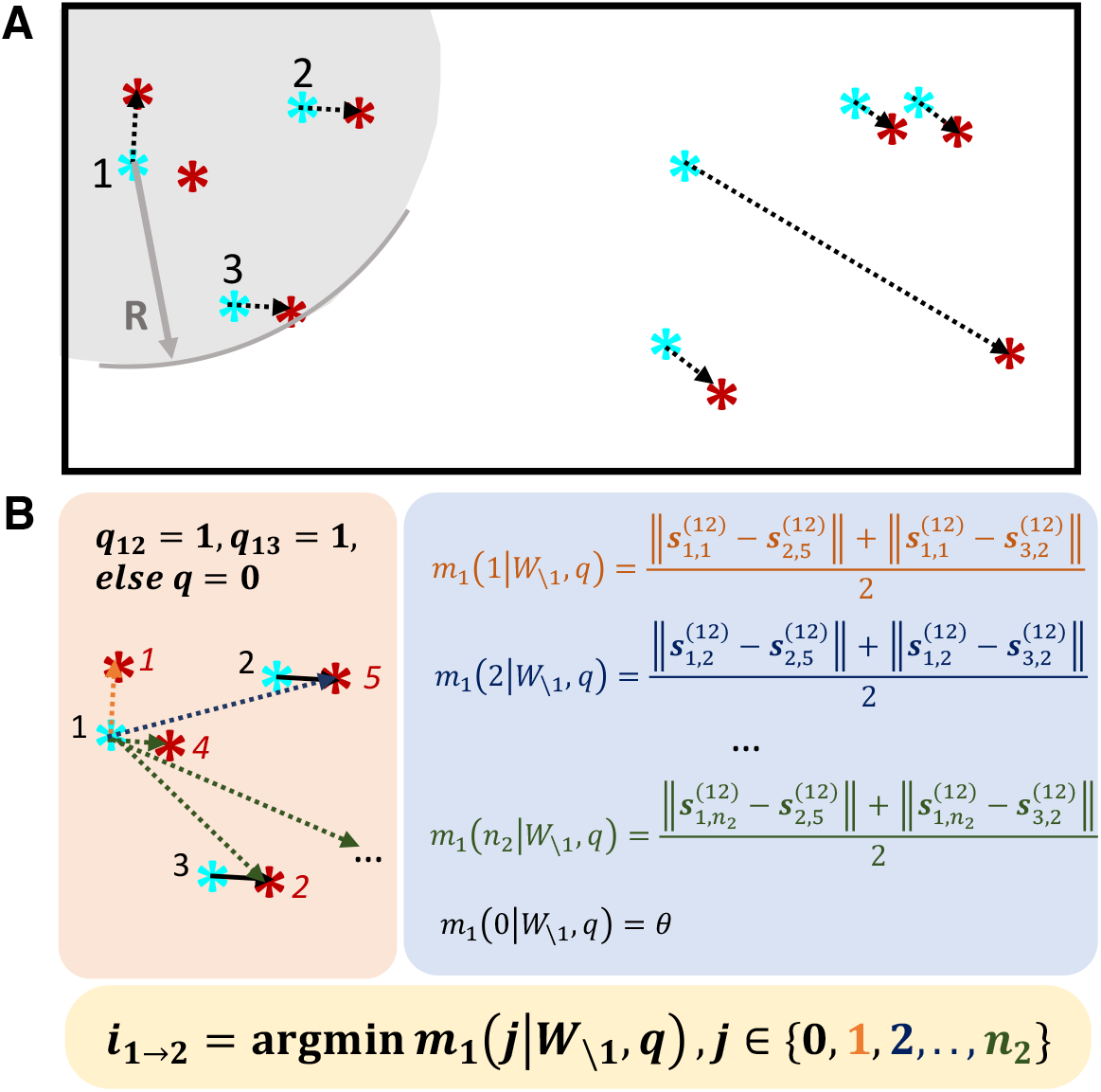
Graphical explanation of the optimization problem

If 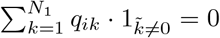, then neuron *i* does not have any neighboring neurons that can provide information. This either means that neuron *i* does not have neighboring neurons or that the neighboring neurons *k* that could have provided information (*q*_*ik*_ = 0) to neuron *i* were all unassigned with neurons in session 2. In this case, we define *m* as

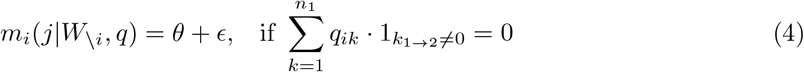

In above, *θ* is a threshold for constraining the maximum motion discrepancy. It is a user defined parameter, in unit of pixel(s) or voxel(s) in the 3D case. The smaller it is, the stricter the matching criterion it will be. *ϵ* is an arbitrary small number. This way of defining this special case shall become clear after we introduce the last special case *j* = 0 where neuron *i* is not matched to any neurons, and after we introduce the optimization objective function in the next section.

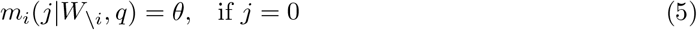

Thus, our full definition of the **discrepancy cost** *m* is:

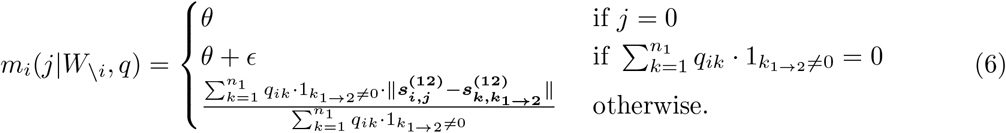

Essentially, this way of defining the special cases will push the case 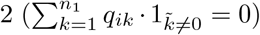 into being the case 1 (*j* = 0)—leaving neurons with no neighbors unassigned during optimization. We introduce the full optimization process below.

### 4.2 The Optimization Problem

We formulate the combinatorial optimization problem as finding a *W* that minimizes the total motion discrepancy of all neurons in session 1,

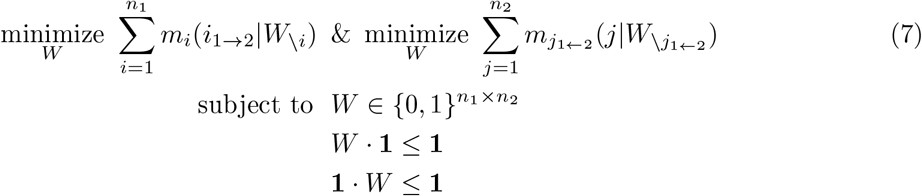

Alternatively, we can sum to only one direction. This objective function is more basic than the one above (equation 7) to solve. In the following, we first introduce the method to solve this reduced objective function.

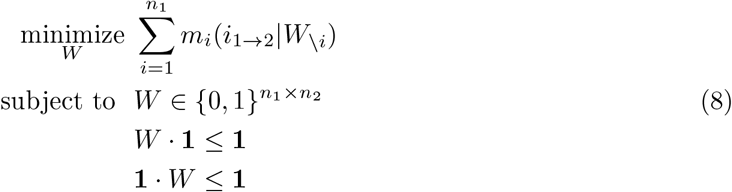

From the definition of *m*_*i*_(*j*|*W*_\*i*_) in equation 6, we can see that the value of one assignment effects other neighboring neurons’ assignments and their assignments in turn will affect the assignment in consideration. Thus, the minimization of the objective function (8) requires us to optimize all pairs of assignments jointly. To the best of our knowledge, there is no known algorithm for solving it within polynomial time.

Here we developed a fast algorithm to solve this optimization heuristically. The idea is to update the assignment of neurons one by one through the previous assignment *W*^*t*−1^ by letting

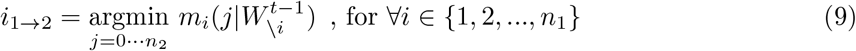

Let’s continue using figure 3 as a concrete example of equation 9. In (B), the blue box calculates all the *m*_1_(*j*) for *j* = 0, 1, 2, …, *n*_2_ based on *W* ^0^. In the yellow box below, we pick one assignment (*i*_1→2_) that gives the minimum *m*_1_. (If *i*_1→2_ = 0, then there is no neuron in session 2 that is suitable for the neuron 1 in session 1.) Repeat this process for all the neurons in session 1, we obtain an updated (*i, i*_1→2_), or *W*^*t*^.

Then, the process repeats from *W*^*t*^ to *W*^*t*+1^; for all the neurons in session 1 we re-calculate *m* given *W*^*t*^ by applying equation 9 again. The iteration procedure goes on until the assignment is stable (*W*^*t*^ = *W*^*t*−1^).

Then, we can run the same algorithm again, albeit in the other direction to obtain the result of the second half of the objective function (7).

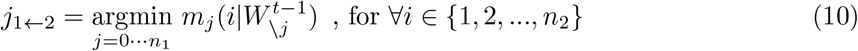

Finally, an intersection of the answers to the equation 9 and 10 gives the approximate optimal *W*^∗^. Note that in each direction, each neuron updates their assignment independently based on *W*^*t*−1^, therefore there could be one neuron given more than one assignment. However, once we take the intersection of the result from updating in either direction, these multi-match cases would not show up.

Algorithm (1) describes the basic matching algorithm for the objective 7 and we will discuss the details of model initialization and manual interventions in the following subsections.

#### Algorithm 1

*STAT* algorithm for matching neurons in two sessions

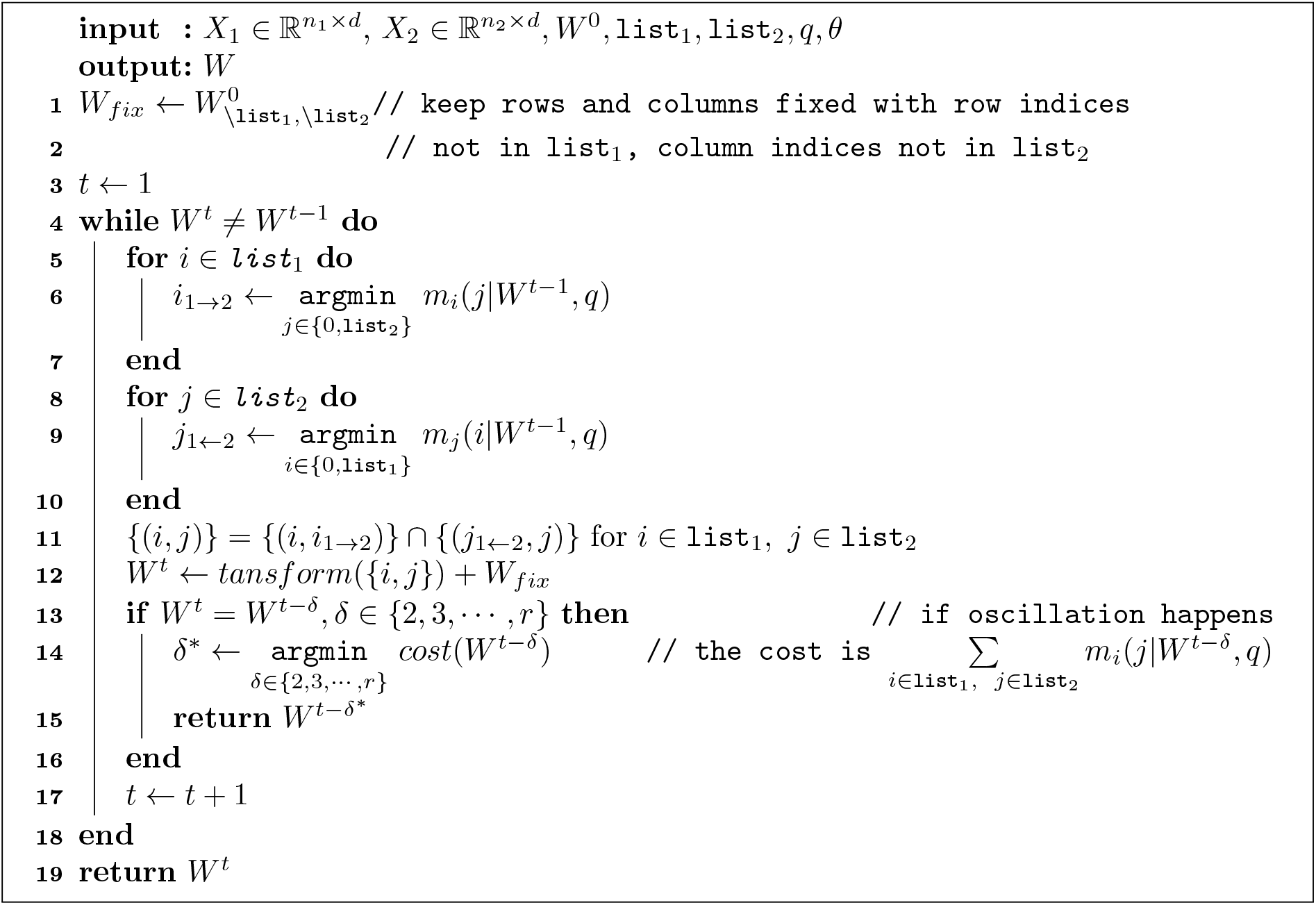

### 4.3 Initialization

Since the initial *W* ^0^ provides some anchoring points for roughly determining the motion vectors, it is important to get at least a few pairs accurate in a small neighborhood. Fortunately, the initialization does not require many pairs of (*i, i*_1→2_) and we can easily get the initialization using different algorithms since *W* ^0^ does not need to be all correct.

In practice, we use the cell shape correlation (see Section 1) to initialize *W*, which is similar to CaImAn [2]. We formulate it as the following problem:

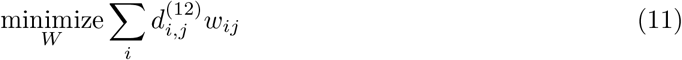

where *W* is a complete matching from session 1 to session 2 if *n*_1_ ≤ *n*_2_ or from session 2 to session 1 if *n*_2_ ≤ *n*_1_; *w*_*ij*_ is used for selecting the neuron in session 2 to match neuron i from session 1. *d*_*i,j*_ could be pairwise shape correlation or pairwise distance 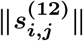.

We use the Hungarian algorithm to solve equation 11, and obtain our initial estimate of *W*, which includes some correct neuron matches and some incorrect neuron matches. We feed this *W* as into equation 6 to start iteration and let the update rule 9 and/or 10 refine *W*.

Alternatively, one can just pick three to five pairs in a small area by hand to initialize.

### 4.4 Manual Intervention

Human checking is helpful as the human visual system is a great outlier detector. Specifically, we color-code each neuron assignment pair by its shape correlation or some other similarity measure. A bad match will tend to have bad shape correlation, and will ‘pop out’ as a potential mismatch. This will be shown in practice in the application of the algorithm in our songbird data in another subsequent section.

## 5 Matching More Than Two Sessions

The previous discussion has been about solving the assignment problem between two sets of neurons. If we are provided with multiple sets of neurons, how can extend our method? Do we need additional theoretical exploration?

Despite that there could be additional theoretical development here, we do not pursue it in this project as we can simply run all sets of neurons in a pair-wise manner, especially because our ***STAT*** algorithm is fast. In all the pair-wise assignment results: (1) there can be an assignment conflict, for example, when session 1 and 3’s matching is different from session 1 and session 2’s matching followed by session 2 to session 3’s matching. If this happens, then any neurons involved in the conflict would be put into a class of “undecided neurons” in the result, and would require manual input to resolve the conflicts; (2) there are neurons without any conflicts; they are considered very confident. From the tests on ***STAT*** so far, as expected, the size of the class “undecided neurons” is related to the total number of pairwise session tracking. At the same time, because the possibility of each individual neuron extracted on all the sessions gets lower as there are more sessions, the class “undecided neurons” is usually not big. In our dataset of neurons from 6 days, each with 60-80 neurons, we only see 1-6 pairs of “undecided neurons” and thus conflicts can be resolved quickly. Users can also apply pairwise session tracking only to adjacent sessions.

## 6 Validation with Simulated Data

In this section we test ***STAT*** on simulated data under different parameters.

### 6.1 2D Data

#### 6.1.1 Generating 2D Data

We first use spatial warping and randomly delete neurons to simulate motion and unique neurons, similar to the real data shown in figure 1) There are many ways we could simulate the warping as long as it gives a smooth motion field. Here we use Gaussian Process (GP) to simulate a smooth and nonrigid motion field. The motion field that GP gives has the relationship that the distance between the two points in the field is inversely related to the correlation between the motion vectors at the two points, which should be a good model for simulating the real motion in the brain tissue. We then apply the motion field to real spatial footprints extracted from our data and randomly delete and add neurons in two sessions. The result of simulated data is shown in figure 4, where we display three datasets in the first row of the three panels.

**Figure 4:**
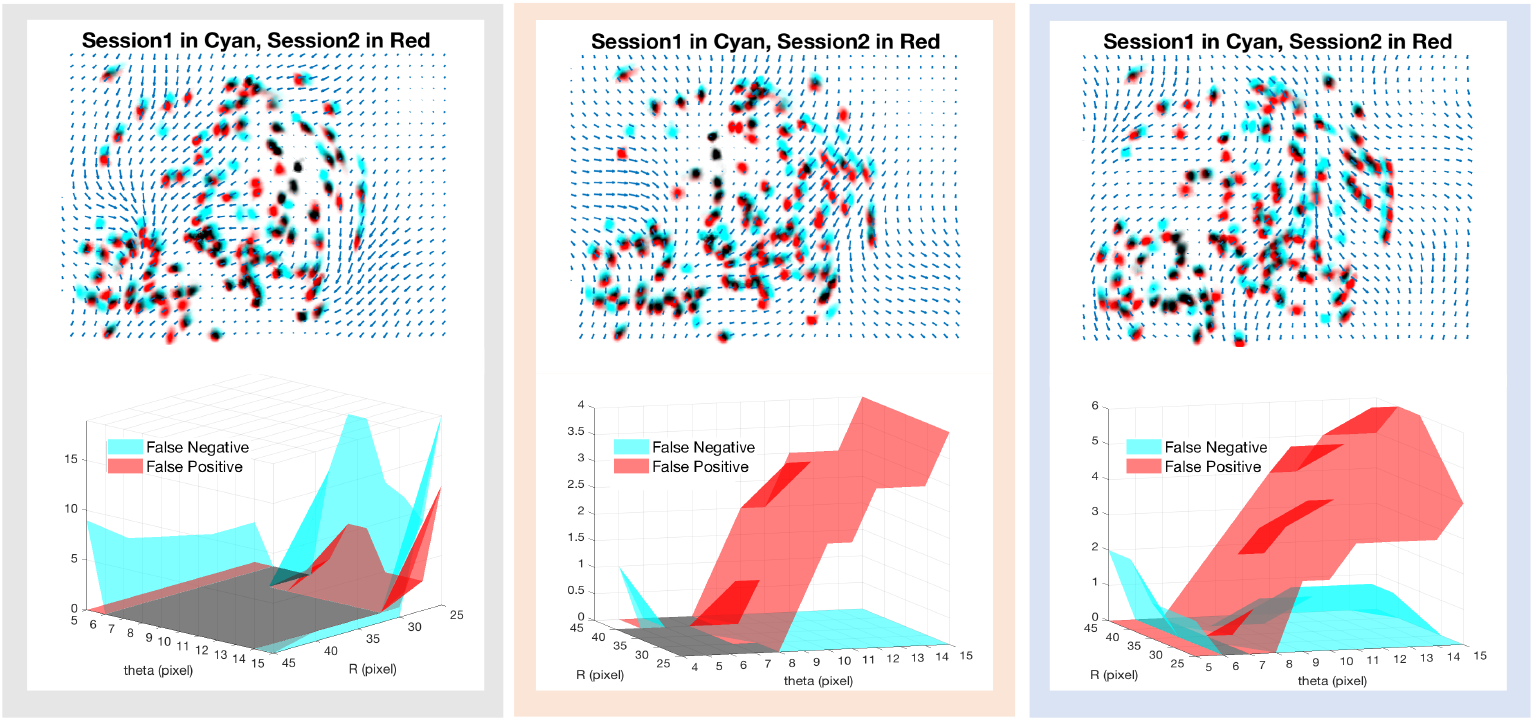
Three 2D simulate data and ***STAT*** result under different parameters.

#### 6.1.2 Testing on Simulated 2D Data and Parameter Characterization

As discussed in the section above, there are two key parameters in the objective function 6. One is the upper threshold of motion discrepancy *θ*, and the other one *q* defines the weights of neighbors in calculating the motion discrepancy. If we choose the simplest step function case in setting *q* (which sets *q*_*ik*_ = 1 if the distance between neuron *i* and neuron *k* is less than or equal to R, and *q*_*ik*_ = 0 otherwise), then we have the two parameters: *θ* and *R*.

Is ***STAT*** sensitive to *θ* and *R*? How should we pick the right parameters for the best performance? Here, we run ***STAT*** on the three datasets we simulate under different combination of the two parameters (see the section above) and compare it to ground truth so that we have the false negative and positive counts as a measure of performance under different parameters. We plot the result in the second row of in the three panels in figure 4. In the figure, we plot false negative count in cyan and false positive count in red, and the region they overlap in a darker color. In some extreme parameter combinations, ***STAT*** fails to converge, in which case there is no data entry in the plot but overall in each dataset there are a large parameter regime of perfect matching.

The three datasets and the results in figure 4 show the general characteristics of the two parameters.

1. Parameter *θ* *θ* is the parameter for controlling how strict our algorithm is: the smaller the *θ* is, the more strict the assignments are, and vice versa. This can be seen in the figure 4, especially in the last two datasets: when *θ* is getting too small, there will be more false negatives; when *θ* is getting too big, there will be more false positives. As to how to pick *θ, θ* should roughly be in the range of visual resolution of the data acquired. For example, in our data, neurons have radius of about 12 pixels, and center points separated by 2-3 pixels are not perceptible. In addition, there are errors of 1-3 pixels when estimating the centers of neurons. Thus, *θ* in our case, would be at least about 5 (pixels). If we take into the account that there is some motion discrepancy among neurons, then we should add in a redundancy of about 2 pixels. We thus have *θ* at about 5-7 (pixels). Indeed, we see that in figure 4, *θ* around these values gives good results while at the same time there can be much sensitivity around the range of 5-7 (pixels).
2. Parameter *R* *R* depends on the neuron size and neurons’ density in the data. Because neurons have radius of about 12-15 pixels in our data, there would not be many neurons within the range of *R ≤* 30 of each neuron so that the cost *m* we calculate would not be informative. Thus, *R* should be large enough to allow some number of neighboring neurons to be included. The first dataset in figure 4 demonstrates this point. At the same time, *R* should not be so big that there are already neurons with drastically different motions included. This side of reasoning is not demonstrated by any of the datasets shown. In general, *R* is less sensitive than *θ*. In general, because ***STAT*** runs fast (time scale of second to minutes), we recommend doing a small parameter sweep across the 2-dimensional parameter space after narrowing down the range of parameters based on the dataset.

### 6.2 3D Data

Due to the large size of volume imaging data, methods for aligning the FOV across sessions are extremely slow. However, in our method, 3D cell tracking is not different from the case in 2D, as we would simply be working with vectors of 1 more dimension. This makes our method suitable for cell tracking in volume imaging.

We still use 2D GP to simulate motion field: we simulate a 2D motion field in each direction and multiply then before taking a cube root to obtain the final 3D motion field. We use this method to simulate 3 motion fields for three datasets, one of which is shown in figure 5(A). We then randomly select 120 points in the volume and apply the motion field to get the second set of neuron centers, before we randomly delete and increase neurons in each set. One simulated dataset is shown in figure 5(B): one set of neuron centers is shown in cyan and the other one is shown in red, with the yellow lines connecting the ones that are ground truth assignments. Comparing it to figure 4, 3D data tends to be harder to visualize.

**Figure 5:**
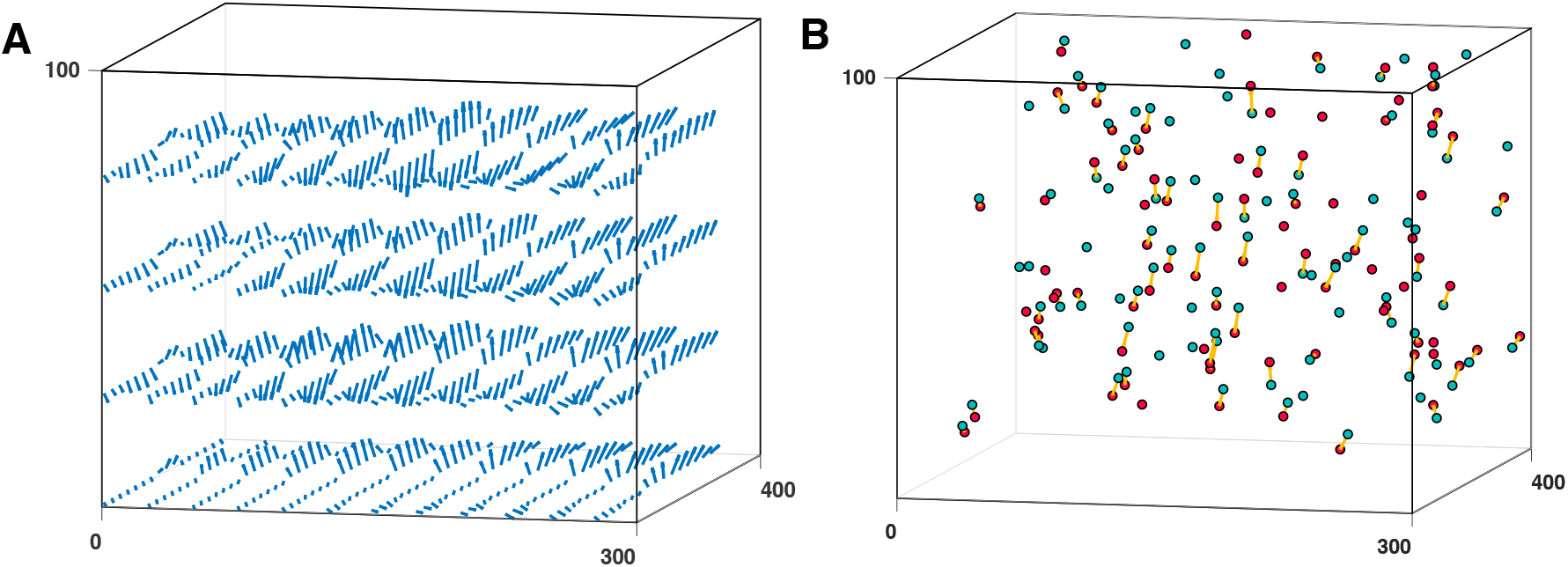
Simulated 3D data. (A) 3D motion field. (B) Simulated neuron centers in two sessions: session 1’s neuron centers is shown in cyan and session 2’s is shown in red, with the yellow lines connecting the ones that are ground truth assignments.

***STAT*** still runs fast and finds us assignments on 3D data. Just as in the 2D case, we did a parameter sweep on three simulated datasets, and collected false negative/positive counts over different combinations of parameters. The result for each of the datasets is shown in the three panels in figure 6. We see that the two parameters’ characteristics are very similar to the 2D case.

**Figure 6:**
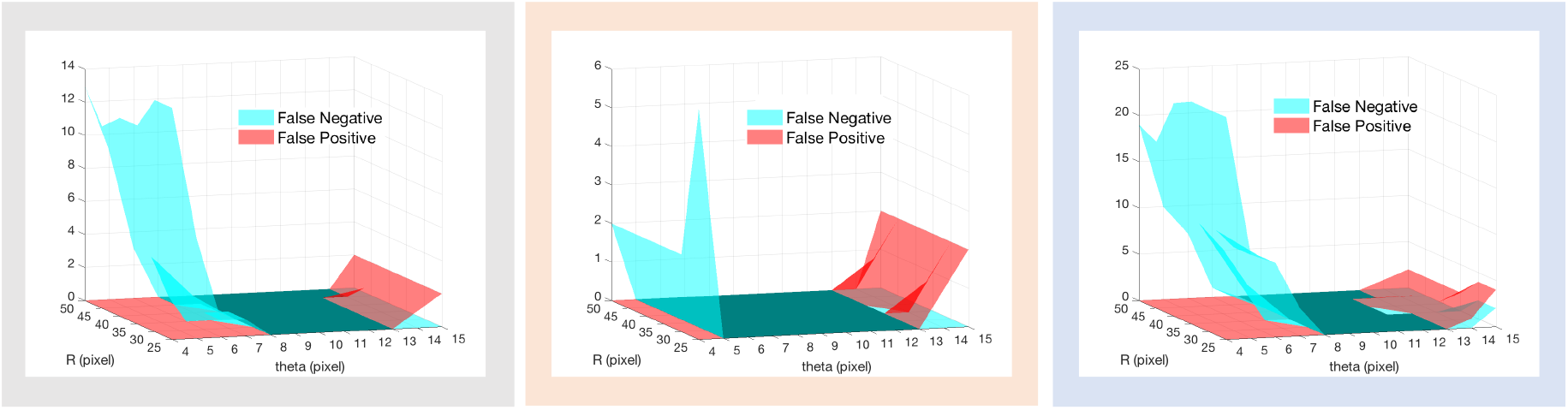
The result of running *STAT* on three simulated 3D data. Graphic schemes are the same as in the second row of figure 4.

## 7 Application to Adult Songbird HVC Recording

This work is motivated by the need to track HVC neurons in late-tutored zebra finches to study how necessary is experience in forming motor sequence. Please refer to [5] for the detailed study. Here we plot 3 figures 7, 8, 9 using data from the study in this document to be self-contained [14].

**Figure 7:**
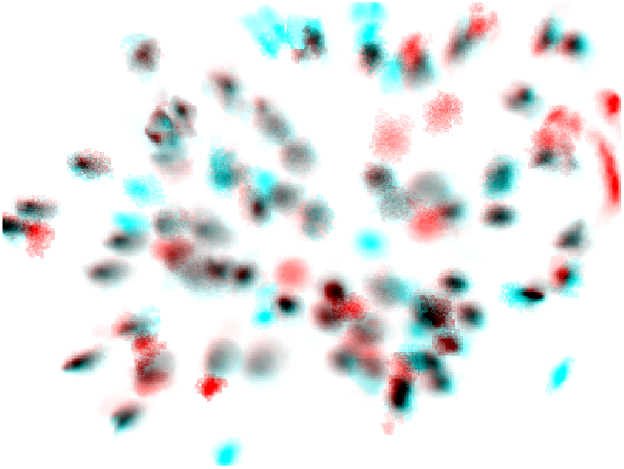
Real data from songbirds. Overlap of neurons from two session of one-photon calcium imaging of HVC brain region of an adult zebra finch: session 1 is in cyan and session 2 is in red. The overlapping regions show darker colors. There is not much motion in the center of the FOV, however neurons on the edge show motion.

**Figure 8:**
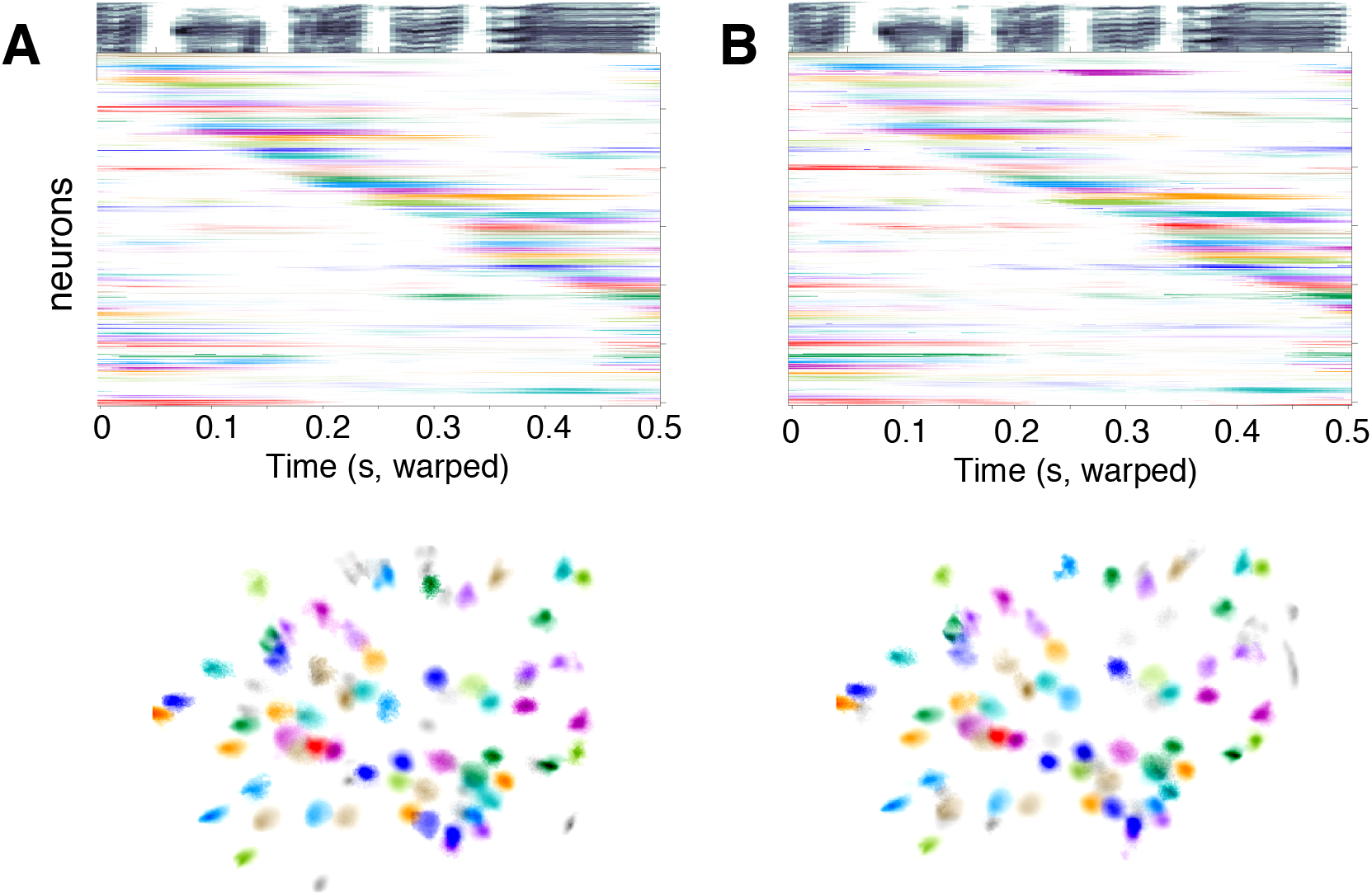
*STAT* ‘s result on the real data shown in figure 7. In the other panel, the same colors other than grey denote an assignment of neurons between the two sessions and color grey denotes the unique neurons within each session. (The colors are randomly assigned for each neuron within session 1.) The upper panel shows the song spectrogram on top in grey scale and the raster below show that the tracked HVC neurons have stable song locked neural activity across the two sessions.

**Figure 9:**
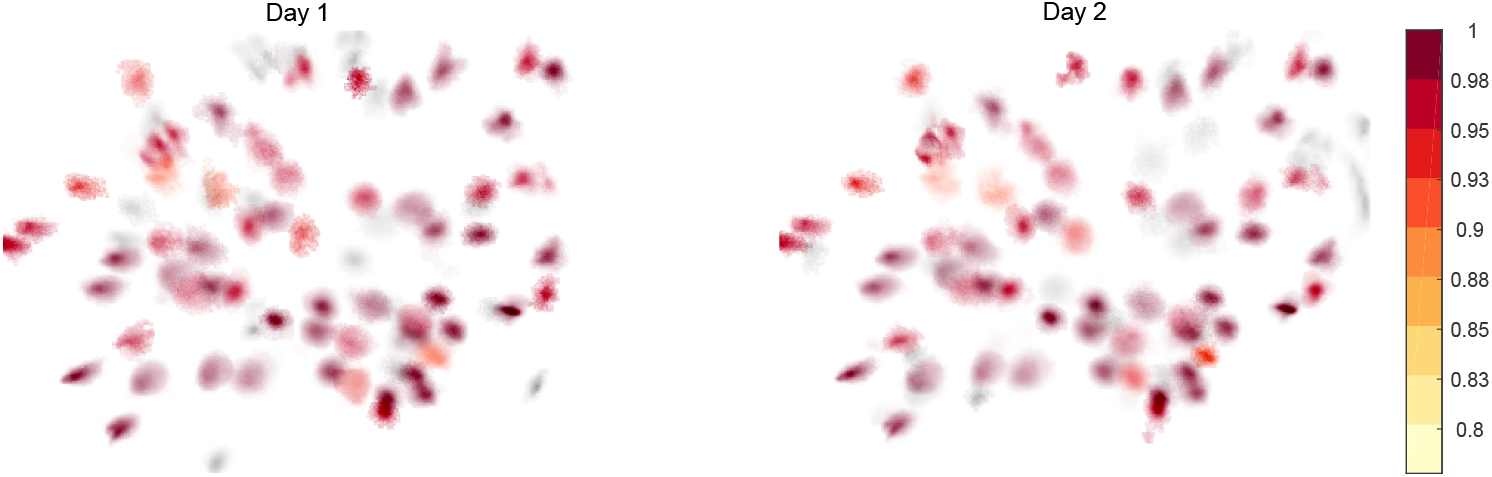
Pairwise shape correlation of assignments. Note that because one-photon imaging is blurry and so the neuron shape information is not reliable as a piece of information in deciding assignments. However, it can be used for sanity check.

## 8 Application on Dense Two-photon data

Here we apply the algorithm on the publicly available two-photon data from the Allen Brain Observatory (ABO) [15]. In addition to tracking the soma, which is a process already included in the ABO pipeline and has results available as metadata, we apply CaImAn[2] to extract dentritic spatial components and apply ***STAT*** to track them along with all the soma components. The result is shown in figure 10 and is quite satisfactory upon visual inspection.

**Figure 10:**
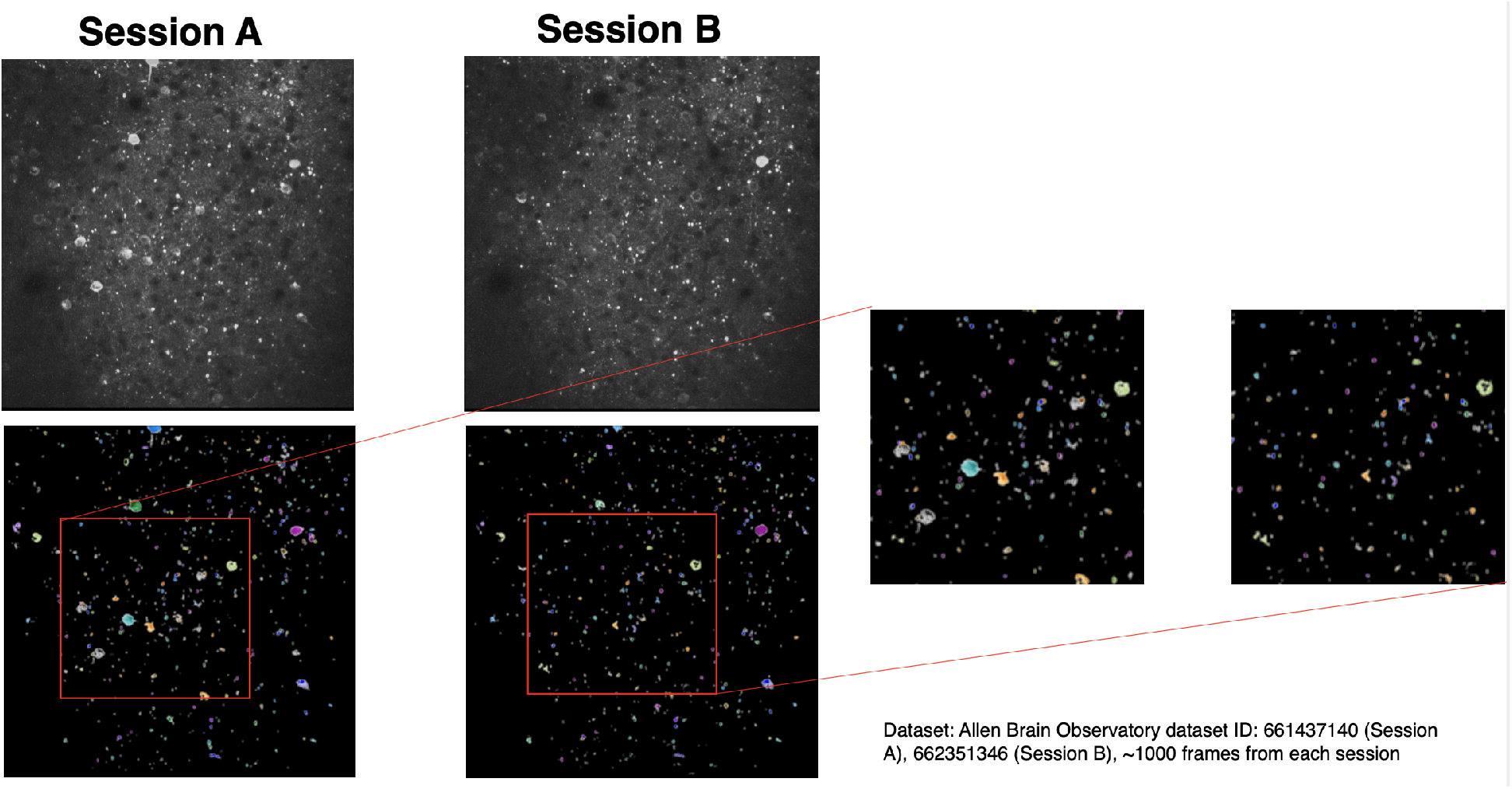
Application on dense two-photon calcium imaging data. Upper panels on the left show the time-averaged image of the two raw videos on Allen Brain Observatory (session ID 661437140 and 662351346 respectively); lower panels on the left show the result of tracking by ***STAT*** where tracked neurons are in the same color across the two session. The zoom-in view gives a more detailed view.

## 9 Discussion

Calcium imaging has been a useful tool in neuroscience, and being able to track neurons in a confident and stable manner is a necessary computational hurdle to overcome to keep up with experimental progress. Our method is meant to do this well. Meanwhile, the problem of tracking neurons can find its root in computer science under the name of other terminologies. Here we first describe how our method relates to some other work in computer science and neurosciences. Then we describe possible extensions.

### 9.1 Similar Work in Computer Science and Neurosciences

Our work in cell tracking is in some sense about finding the transformation (warping) between two images by using a number of neurons as features. Using features to find the transformation between the two images dates back as early as to D’Arcy Thompson’s work On Growth and Form in 1917 (figure 11). From then till today, the problem of finding the warping or spatial transformation has been embedded into various mathematical models and many solutions have been found in various conditions. We introduce a couple perspectives toward the problem below and comment on how our method fits with these perspectives.

**Figure 11:**
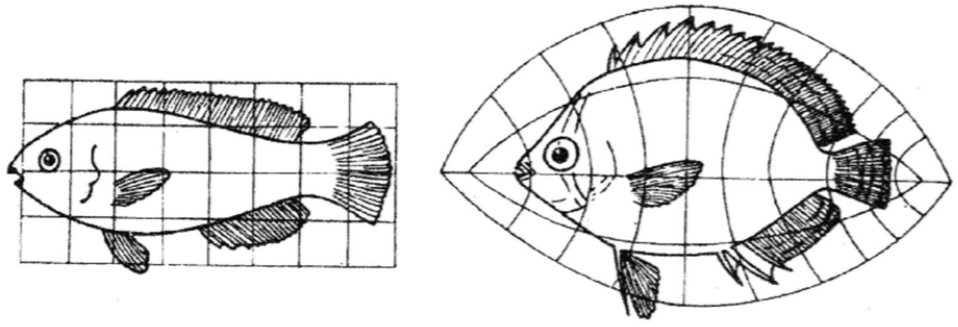
Examples of finding the spatial transformation between two images. This example is in D’Arcy Thompson’s work On Growth and Form in 1917. The figure is copied from reference [16].

In the problem of tracking neurons, if neurons are reduced to centers, just as in our method, then the problem of finding assignments can be viewed as a *point set registration* (PSR) problem. In the many PSR studies, the cost function in [16] is very in line with our method, while the additional smoothness constraint is formulated in a more sophisticated fashion in the various work by [17, 3, 4]. Study [4, 3] use a full probabilistic formulation. Readers are encouraged to try out Ma’s package at https://github.com/jiayi-ma/PR-GLS as well. In practice, our method and the [4] full probabilistic method gives identical results on our songbird dataset upon manual inspection.

In neuroscience, the first paper to the best of our knowledge to have used PSR framework in tracking neurons is reference [18], where a *thin plate spline* (TPS) function is used to model the transformation between the two imaging planes before finding the assignments of neurons. In previous methods without the framework of PSR, such as reference [1, 2], a transformation field was also first estimated before calculating various metrics based on the transformation and finding the exact assignments of neurons. We propose in this manuscript that an iterative optimization of both the general motion field and the exact assignments of neurons simultaneously is a better approach to calcium imaging cell tracking problem. Iterative PSR methods have been used in aligning electron microscopy data [19], two-photon tomography data[20].

### 9.2 Extension: Infer the Spatial Transformation

As noted in references [10, 18], a thin plate spline (TPS) function could model real brain tissue motion well. From the assignments of neurons found by ***STAT***, we can directly obtain the transformation at neuron centers. By interpolating and extrapolating using the TPS model, we can obtain the motion at any point in the field of view. Figure 12 shows in the first row of each panel the ground truth motion field (blue arrows) of the three simulated 2D datasets (same as in figure 4) and in the second row the inferred motion field by fitting the TPS model over the motion vectors at neuron centers. Except for the motion fields on the edges of the field of view (far from neurons), the estimate and the ground truth are very similar.

**Figure 12:**
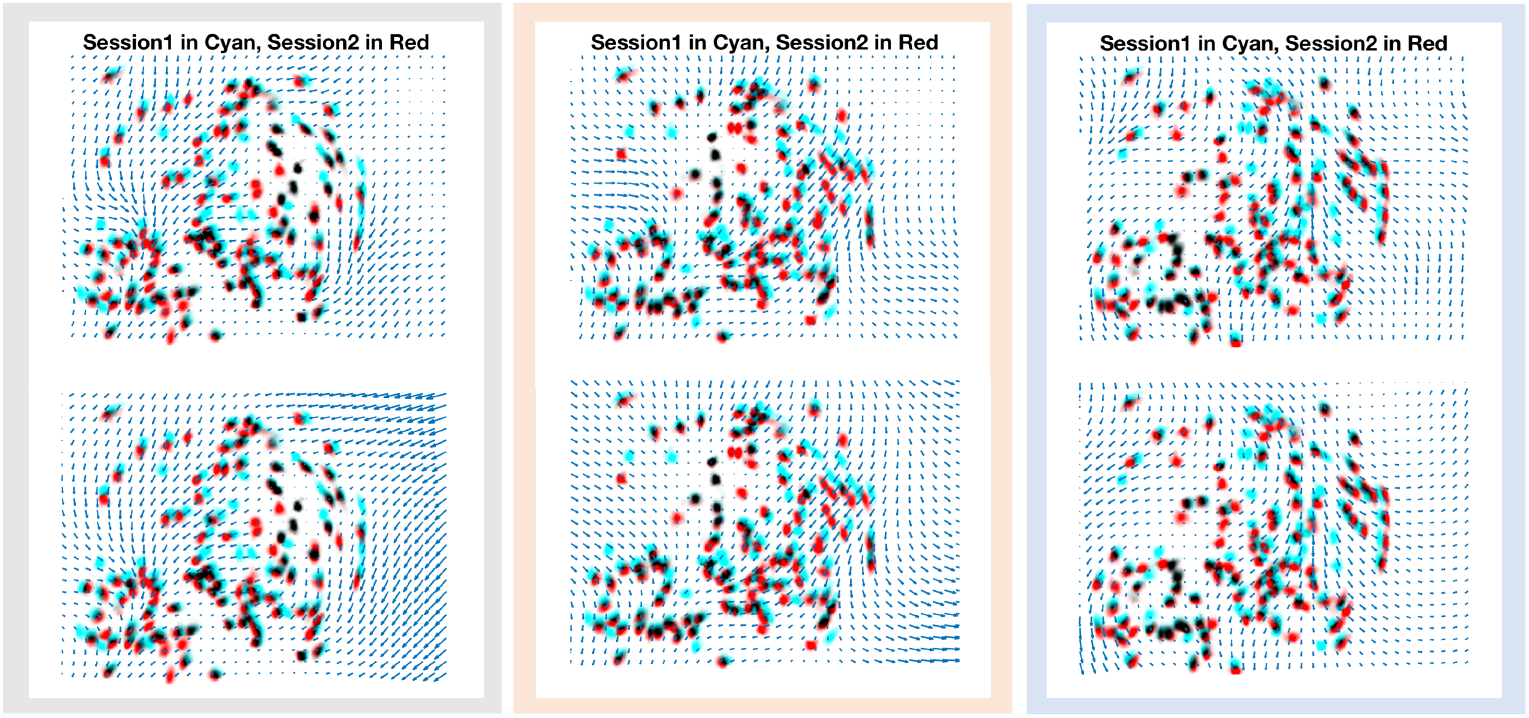
Ground truth motion field and estimated motion field using TPS model. In the three simulated 2D datasets, we here plot the ground truth motion field again in the first row of each panel and in the second row of each panel the estimated motion field for each data using the TPS model from the result of ***STAT***

There are many future directions that could make use of the estimated motion field. Cell extraction algorithms for processing raw calcium imaging videos often has certain thresholds to set real neurons from false positives [9, 13]. Thus, setting a threshold too high for certain sessions could lead to missing neurons in the result of cell extraction of these sessions. Using the inferred spatial transformation, we can infer the location of certain missing neurons and thus potentially re-extract their neural activity. As another use of the inferred spatial transformation and inspired by reference [18], it could potentially be used as a sanity check of the assignment found by ***STAT***.

Specifically, to check if the assignment of neuron *i* and neuron *i*_1→2_ makes sense, we could use all the other assignments to estimate the motion field and infer where neuron *i* could be in session 2. We then compare this estimate to the location of the neuron *i*_1→2_. The error should be small if the assignment is correct and TPS is a reasonable model of the motion field. Usually, however, we find that pairwise correlation of the assignments is a sufficient sanity check (as in figure 9).

## 10.2 Acknowledgements

This work was supported by a grant from the Simons Collaboration for the Global Brain, the National Institutes of Health (NIH) [R01 DC009183] and the G Harold and Leila Y Mathers Charitable Foundation. S. Gu received partial funding from the ShanghaiTech University Undergraduate Study Abroad Scholarship. E.L.M. received support through the NDSEG Fellowship program and the Simons Society of Fellows.

We thank Liam Paninski for his prudent thought in the matter of calcium imaging cell tracking and encouragements throughout the project, Andrew Bahle for early testing the package, and Kail Miller for ideas on illustrations.

## 10.3 Code Availability

The code and sample data is available here at: https://github.com/shijiegu/STAT

